# Programmed Cell Death Modifies Neural Circuits and Tunes Intrinsic Behavior

**DOI:** 10.1101/2023.09.11.557249

**Authors:** Alison Kochersberger, Dongyeop Lee, Mohammad Mahdi Torkashvand, Sandeep Kumar, Saba Baskoylu, Titas Sengupta, Noelle Koonce, Chloe E. Emerson, Nandan V. Patel, Daniel Colón-Ramos, Steven Flavell, Andrew M. Leifer, Vivek Venkatachalam, H. Robert Horvitz, Marc Hammarlund

## Abstract

Programmed cell death (PCD) is a common feature of animal development. During development of the *C. elegans* hermaphrodite, programmed cell death eliminates 131 cells in stereotyped positions in the cell lineage, mostly in neuronal lineages. Blocking cell death results in supernumerary “undead” neurons. We find that undead neurons can be wired into circuits, can display activity, and can modify specific behaviors. The two undead RIM-like neurons participate in the RIM-containing circuit that computes movement. The presence of these two extra neurons results in animals that initiate fewer reversals and lengthens the duration of those reversals that do occur. We describe additional behavioral alterations of cell-death mutants, including in locomotory turning angle and pharyngeal pumping. These findings indicate that physiological or evolutionary variations in PCD might reveal latent neuronal elements that the nervous system can incorporate to modify nervous system function and animal behavior.

## Introduction

The somatic cell lineage of the *C. elegans* hermaphrodite contains 131 programmed cell deaths (PCDs)^1,2^. These cell deaths are a feature of normal development, unlike cell deaths induced by damage or disease. PCD occurs during the development of most and possibly all animals^3–5^. Of the PCDs that occur during *C. elegans* hermaphrodite development, 105/131 (80%) are “neural-proximate,” i.e. derive from branches of the cell lineage that generate only neurons and neuron-associated glial-like cells^1,2^. 94/131 (72%) of the cell deaths occur in lineages that give rise exclusively to neurons. By comparison, only 302/959 (31%) somatic cells in the mature hermaphrodite are neurons (Figure S1A–C)^1,2,6,7^. Thus, PCD is concentrated in neuronal lineages during *C. elegans* development, indicating that PCD plays a key role in shaping the nervous system of this animal.

Study of the fates and functions of cells normally eliminated by PCD is possible using mutations that disrupt the core mechanisms of cell death. For example, *C. elegans* mutants deficient in the function of the gene *ced-3* lack essentially all somatic PCDs because they are defective in the major caspase that drives PCD^8,9^. *ced-4* mutants lack all somatic PCDs because they cannot activate *ced-3*^8,10–13^. In such *ced-3* and *ced-4* mutants, the 131 cells that normally are removed by PCD survive and are referred to as “undead”^14^. Many undead cells become neuron-like^6,8,14,15^. For example, the two serotonergic NSM neurons each have a sister that is normally removed by PCD. In cell-death mutants, the undead NSM sisters express serotonin and morphologically resemble the NSMs^8,15^. Studies of the undead sister of the M4 neuron^6^, undead sisters of the two PVD neurons^16^, undead lineal homologs of the VC neurons in the ventral nerve cord^17^, and other undead cells similarly indicate that undead cells in neuronal lineages share characteristics with normal neurons, including acquiring neuronal morphologies and expressing neurotransmitters. We found that undead neurons often expressed fluorescent markers normally specific to neuronal relatives, suggesting similar patterns of gene expression and revealing similar morphologies of undead cells to those of their normal neuronal relatives (Figure S1D–L). Thus, in the absence of cell death, the *C. elegans* nervous system contains a large number of supernumerary undead neuron-like cells. These considerations indicate that if all undead cells in neuronal-specific lineages acquire a neuronal fate, hermaphrodites that lack PCD have at least 94 extra neuron-like cells, in addition to their normal complement of 302 neurons.

Do undead neuron-like cells function? In *C. elegans*, one undead cell can partially compensate for the function of its sister neuron (M4) after that sister is killed by laser microsurgery^6^. Recent studies of *Drosophila* have similarly found that blocking PCD in specific neuronal lineages produces additional neuron-like cells that can induce a behavioral response when stimulated in decapitated flies^18,19^. These data raise the possibility that undead neuron-like cells are functional neurons that can affect behavior not only in animals with portions of their nervous systems removed but also in animals with an intact, complete nervous system. However, *C. elegans* cell-death mutants appear to be grossly normal in locomotion, egg laying and chemotaxis, raising the question of what function — if any — PCD serves in modulating the activity of the *C. elegans* nervous system^8^.

## Results

### Undead RIM sisters form synapses onto wild-type circuits

To examine the function of undead neuron-like cells in behavior, we studied the sister cells of the two RIM neurons, a bilaterally symmetric pair of interneurons that are integral components of reversal circuits in *C. elegans* locomotion^20–22^. Each RIM has a sister cell that is normally removed by PCD (Figure 1A). To examine the fate of these RIM sisters, we generated a RIM-specific GFP marker, pRIM::GFP^23^ (see Methods). Nearly all animals carrying a canonical and presumptive null *ced-3(n717)* allele^8^ had 4 cells expressing pRIM::GFP rather than 2 in wild type, indicating that the undead RIM sisters typically adopt a neuronal, RIM-like fate (Figure 1A). By contrast, other undead neurons often did not express terminal fate markers of their sister (Figure 1B-G). In some cases, we observed *ced-3* mutants with more cells expressing the marker than expected. This may be a consequence of undead cells dividing after adopting their neuronal fate; alternatively, undead cells in other lineages may have mixed fates and so may express markers that are not expressed in their sister (Figure 1B-G).

**Figure 1.**
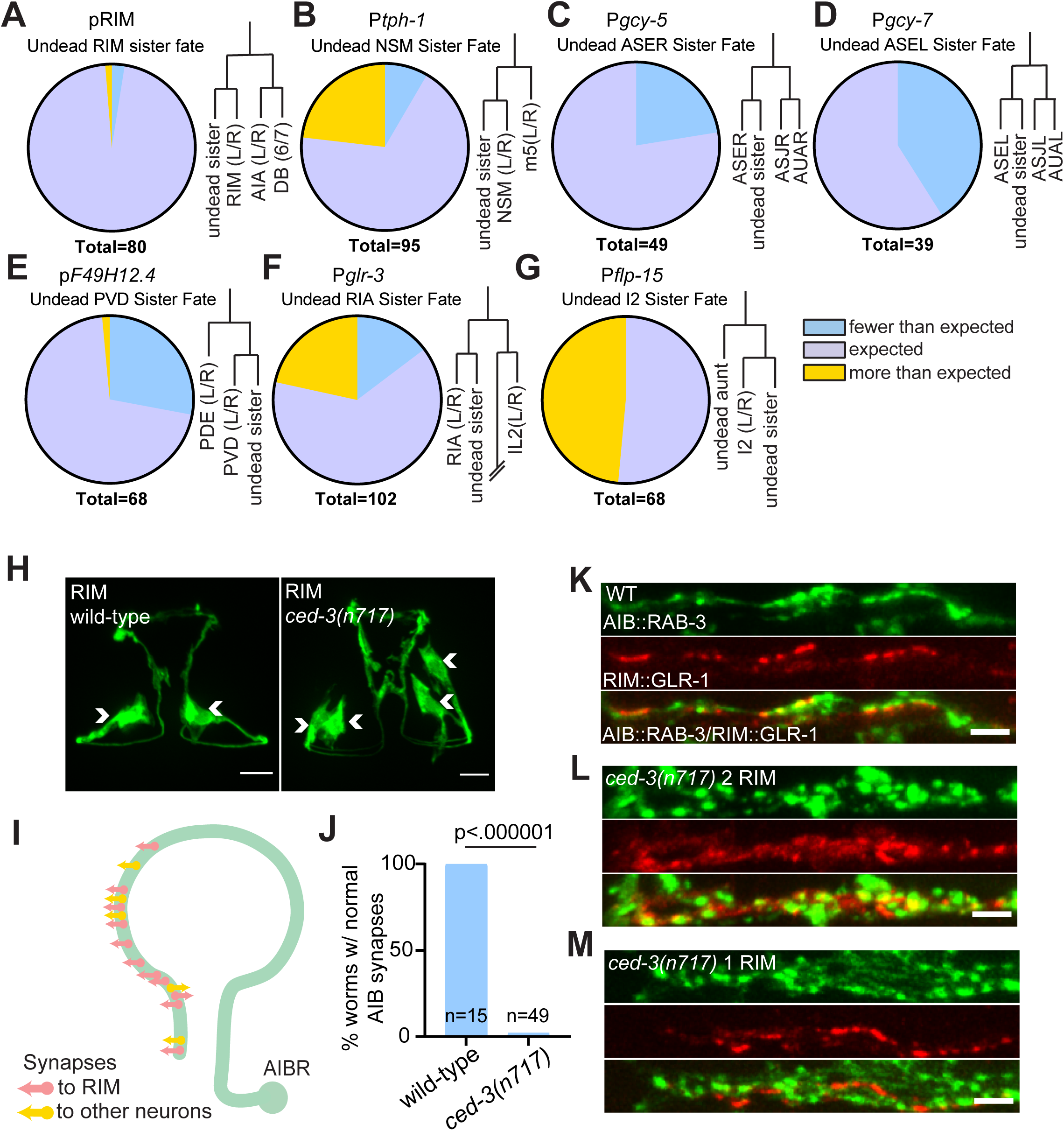
Undead RIM sister cell form synapses with AIB, altering presynaptic patterning. (**A-G**) Fate marker analysis in the *ced-3(n717)* (cell-death mutant) background of selected neuron types with undead relatives. Each panel shows the percentage of animals exhibiting fewer or more than the ‘expected number’ of cells, as well as a representation of the local cell lineage^1,2^. The ‘expected number’ is defined as the number of neurons of the type present in wild-type animals, plus the number of lineage-proximate undead cells. The following markers were used for each cell type (also see Methods and Table S3): RIM: pRIM; RIA: *glr-3*; I2: *flp-15*; PVD: F49H12.4; ASEL: *gcy-7*; ASER: *gcy-5*; NSM: *tph-1.* (**H**) Micrograph of RIM fate marker in wild-type and *ced-3(n717)* (cell-death mutant) backgrounds. Arrowheads indicate cell bodies. The wild-type animal has two RIM neurons, and the *ced-3(n717)* animal has two additional RIM-like cells and is counted as contributing to the orange “more than expected” sector in panel (A). 10 µm scale bars. (**I**) Schematic of AIB synaptic connections to RIM and other neurons^24^. (**J**) Quantification of animals with stereotyped AIB synaptic patterning in wild-type and *ced-3(n717)* animals. (**K**) RAB-3::GFP presynapses in AIB (top panel) and GLR-1::tagRFP postsynapses in RIM (middle panel) colocalize (bottom panel). (**L**) *ced-3(n717)* mutants display an increase in AIB presynapses (top panel) and RIM postsynapses (middle panel) when both RIM and RIM-like cells are labeled. (**M**) *ced-3(n717)* mutants display an increase in AIB presynapses (top panel) but display normal RIM postsynapses (middle panel) when only one of the RIM and RIM-like cells is labeled because of mosaicism of the labeling array. 2 µm scale bars. Statistical significance was determined by Fisher’s exact test.

All 4 RIM-like neurons had similar morphologies, indicating that they underwent similar developmental programs in addition to expressing a terminal fate marker (Figure 1H). In the normal nervous system, each RIM receives synaptic input from one of the bilaterally symmetric AIB sensory neurons. Although AIB synapses onto other neurons, RIM is its primary postsynaptic target (Figure 1I)^24^. Synaptic inputs from AIB to RIM are received by the postsynaptic glutamate receptor GLR-1^22,25^. Unlike the RIM neurons, the AIB neurons have no lineage-proximate PCD events. Thus, the AIB-RIM circuit provides a system for investigating the connectivity and function of undead neuron-like cells.

We visualized synapses from AIB onto RIM using RAB-3::GFP to mark presynaptic regions in the AIBs and GLR-1::tagRFP to mark postsynaptic regions in the RIMs and undead RIM sisters. We found that *ced-3(n717)* animals have a completely penetrant patterning defect at AIB-RIM synapses (Figure 1J). In control animals, both the presynaptic RAB-3 marker (in AIB) and the postsynaptic GLR-1 marker (in RIM) had a punctate organization. Pre- and postsynaptic puncta were often apposed to one another, consistent with these markers localizing at sites of synaptic transmission (Figure 1K). *ced-3* mutant animals had more numerous postsynaptic puncta in RIM-like cells (the RIMs plus their undead sisters) and more numerous presynaptic puncta in the AIBs (Figure 1L). We hypothesized that this observation indicated that in *ced-3* mutants the AIBs form an increased number of synapses: one set of synapses onto the normal RIMs and another set onto undead RIM-like neurons, with the axons being too close together to resolve by imaging.

To better understand the nature of AIB-RIM synapses in *ced-3* mutants, we performed genetic mosaic analysis. Because the transgene carrying the pre- and postsynaptic markers is lost at low frequency during mitotic divisions, we could identify animals that carried the transgene in AIB and in only one of the presumptive RIM/undead RIM pair. In these animals, the distribution of GLR-1 in the RIM or undead RIM was largely normal (Figure 1M, center). These data indicate that the increase in GLR-1 signal observed in RIM-like cells in *ced-3* animals (Figure 1L, center) involves contributions from both the normal and undead RIMs. Furthermore, in these animals with only a single labeled RIM the increases in presynapses in AIB (Figure 1M, top) were similar to those in animals with 2 RIM-like cells in which both normal and undead RIMs were marked (Figure 1L, top), indicating that the increase in AIB synapses is induced by the presence of the undead RIM neuron itself, rather than being a marker-dependent effect. Together, these results confirm that the presence of undead neurons results in wiring changes in the *C. elegans* nervous system^6,14^. Specifically, these data indicate that AIB is synaptically connected to both the normal and the undead RIM neurons in animals that lack PCD, altering the architecture of this circuit.

### Undead RIM sisters are functional neurons that alter behavior

A primary function of neuronal circuits is to control behavior. The AIB and RIM neurons are key components of a well-characterized neuronal circuit that regulates locomotory reversals, an important behavioral activity computed by the *C. elegans* nervous system^21,26–31^. To determine whether the undead RIM-like neurons participate in the reversal circuit, we examined activity in both the normal and the undead RIM neurons using an experimental setup that allowed us to ascertain correlation between activity and reversal behavior. Activity was assessed using the calcium indicator GCaMP6s, which we expressed in both the normal RIM and the undead RIM-like neurons under the pRIM promoter (see Methods) (Figure 2A). We used an anatomical criterion to distinguish normal from undead RIM neurons in *ced-3* mutants: of neurons expressing GCaMP6s, anterior neurons were assigned as the undead RIM sisters, as the anterior daughter of the RIM neuroblast normally undergoes programmed cell death. We confirmed this assignment by observing that these anterior cells were labeled by a *Pegl-1*::GFP::H2B marker -- *egl-1* is a proapoptotic gene expressed in cells targeted for PCD^32,33^ (Methods). A dual-camera system was used to acquire simultaneous images of GCaMP and tagRFP, to adjust for photobleaching and artifacts (Figure S2A). To enable dual observation of both GCaMP6s signals and behavior, we developed an immobilization strategy that preserves high-quality imaging while allowing animals to display forward and reverse movements, although the frequency and duration of these movements differ from freely behaving animals (Figure S2A; Methods).

**Figure 2.**
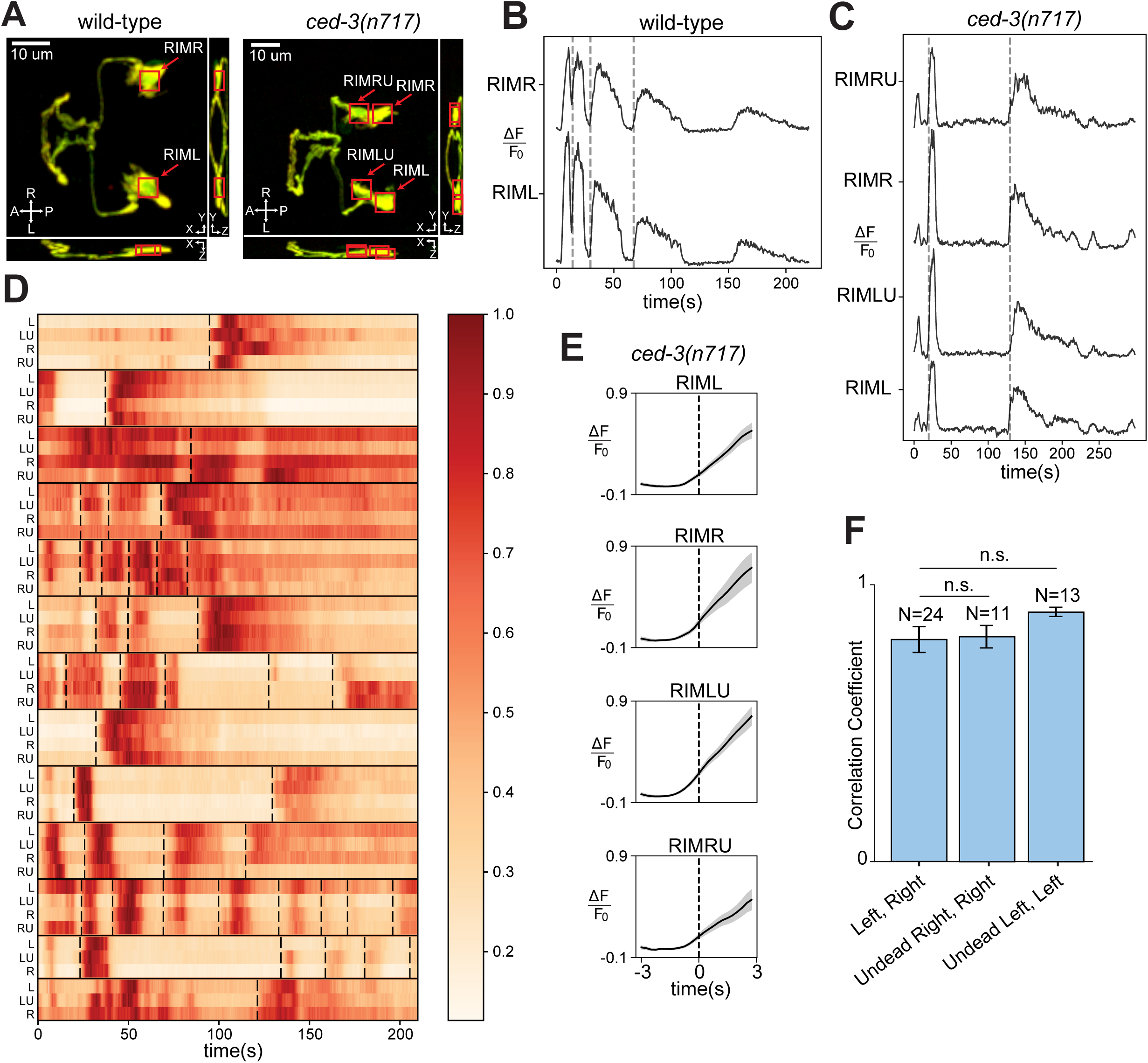
Undead RIMs are functional and their activity corresponds to reversal behavior. (**A**) Annotated micrographs of RIM neurons in wild-type and *ced-3(n717)* animals labeled with GCaMP6s and tagRFP. Red boxes show the cell-body regions that were analyzed. Four cells are boxed for *ced-3(n717)* animals: the two RIM neurons (RIML and RIMR) and the two undead RIM-like cells (RIMLU and RIMRU). (**B**) Calcium traces of RIM neurons in a wild-type animal. Dashed vertical lines represent reversal events. (**C**) Calcium traces of RIM and RIM-like neurons in a *ced-3(n717)* animal. Dashed vertical lines mark reversal events. (**D**) Heatmap of calcium traces of RIM and RIM-like neurons across all *ced-3(n717)* animals. Each row corresponds to a RIM (L, R) or RIM-like (LU, RU) neuron, with its name on the left side of the row. Individual worms are grouped within horizontal black lines. Dashed vertical lines mark reversal events. (**E**) Averaged calcium traces in specific neurons in *ced-3(n717)* animals, aligned on reversal events (dashed vertical lines). (**F**) Correlation coefficients between different pairs of specific neurons in *ced-3(n717)* animals.

Consistent with previous results^27,34^, we found that in control animals RIM activity was correlated with reversals. Specifically, in both in the left and right RIM neurons a sharp increase in GCaMP fluorescence was observed close to the onset of each reversal (Figure 2B). Next, we examined *ced-3* mutant animals that expressed GCaMP6s in two additional RIM-like neurons. We found that undead RIM-like neurons displayed GCaMP signals like those of their normal sisters, typically a sharp increase in fluorescence corresponding to the onset of each reversal (Figure 2C and S2B). We were able to image all four RIM and RIM-like neurons in 11 *ced-3* animals, and three neurons in two additional *ced-3* animals in which we could not resolve the fourth neuron. Across this data set, every animal displayed obvious correlation between overall activity and reversals and also displayed correlation among the activities of all individual analyzed cells (Figure 2D). By contrast, the length of time between two reversals and the pattern of calcium activity after each reversal differed among events (Figure 2D). To further analyze the relationship between the GCaMP signal in individual neurons and behavior, we aligned signals from each individual neuron type according to reversal (Figure S2B; Methods) and averaged across animals. This analysis confirmed that each individual neuron type (the left and right normal RIMs as well as the left and right undead RIMs) displayed activity roughly coincident with—and slightly preceding—reversal initiation (Figure 2E). To determine the amount of correlated activity between these different types of neurons, we calculated the Pearson correlation between different neurons within each animal. We found that the average correlation between the normal left and right RIM neurons was not statistically different from the correlation between each normal RIM neuron and its undead sister (Figure 2F). Overall, these data indicate that in *ced-3* mutants the undead RIM sisters receive functional synaptic inputs that produce activity that is indistinguishable from that of their normal counterparts.

Does the activity of these extra RIM-like neurons influence behavior? Since RIM activity correlates with reversals, we quantified the number of reversals initiated in freely behaving worms. We found that *ced-3* mutant worms initiated ∼60% fewer reversals than wild-type worms (wild type = 2.7 reversals/min; *ced-3* = 1.1 reversals/min; p < 0.0001). In addition, reversals by *ced-3* mutants were of longer duration than those of wild-type worms (wild type = 2.45 seconds; *ced-3* = 3.80 seconds; p = 0.0002). A similar trend is seen for *ced-4* mutants, in which PCD is also prevented (Figure 3A and 3B). These behavioral changes are consistent with the hypothesis that coordinated activity of the extra RIM-like neurons in *ced-3* animals alters reversal behavior. However, *ced-3* mutant animals have many supernumerary neuron-like cells in addition to extra RIM-like neurons that might also influence reversal behavior (Figure S1).

**Figure 3.**
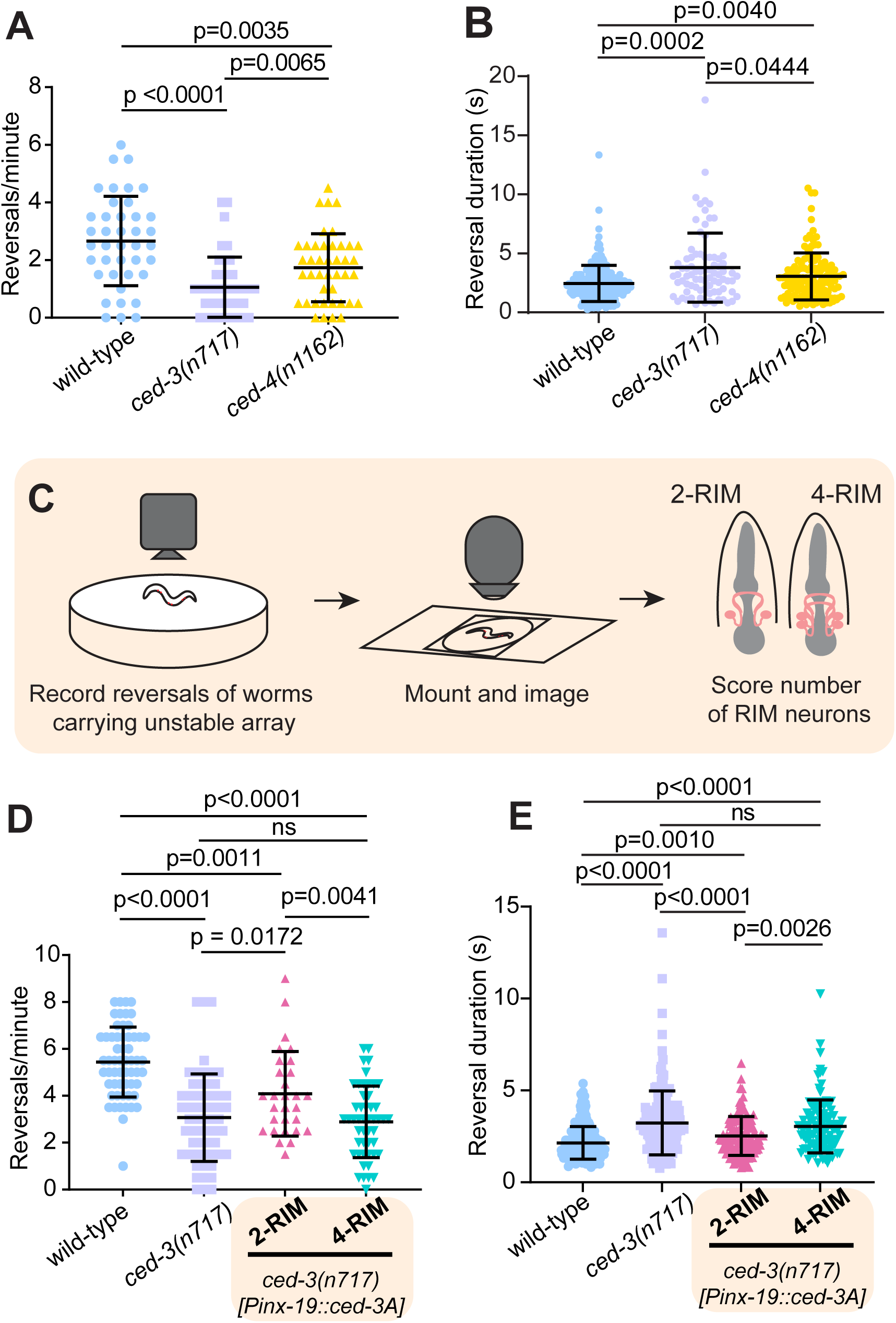
Undead RIMs alter reversal frequency and duration. (**A**) *ced-3(n717)* and *ced-4(n1162)* animals initiate fewer reversals than wild-type animals. Data are mean ± S.D. with individual data points shown. (wild-type n=41, *ced-3(n717)* n=42, *ced-4(n1162)* n=42) (**B**) Reversal durations increased in *ced-3(n717)* and *ced-4(n1162)* animals. Reversal times measured from events in Fig 3A. (**C**) Schematic of behavioral experiment with RIM-specific cell-death rescue. (**D**) *ced-3(n717)* animals initiate fewer reversals than wild-type animals, and removal of additional RIMs partially rescues reversals. Data are mean ± S.D. with individual data points shown. Wild-type, n=58; *ced-3(n717),* n=58; *Pinx-19::ced-3A* (4-RIM), n=49; *Pinx-19::ced-3A* (2-RIM), n=29. (**E**) Reversal length is increased in *ced-3(n717)* animals, and this increase is partially rescued by RIM-specific re-introduction of functional CED-3A using the *inx-19* promoter. Data are mean ± S.D. with individual data points shown. Reversal times measured from events in Fig 3D. Statistical significance was determined by Welch’s t-test.

To determine if the behavioral difference in reversals of *ced-3* animals is specifically caused by the presence of extra RIM-like neurons, we performed mosaic analysis of the RIM cell lineage. We generated a strain of *ced-3* mutants that carries a transgene expressing a functional *ced-*3 cDNA expressed in RIM (and some other neuron types, Methods) under the *inx-19* promoter^35,36^. This transgene is lost at low frequency during mitotic cell divisions (Methods).

Thus, this strain produces animals that have either no undead RIMs (because the transgene is retained in both the left and right RIM lineages, restoring cell death), one undead RIM (transgene is lost in either the left or the right RIM lineage), or two undead RIMs (because the transgene is lost in both left and right RIM lineages, although it is still present in other cells).

These animals also carry a separate stable fluorescent marker for the RIM and undead RIM neurons, *pRIM::tagRFP*, allowing the number of RIM-like cells to be determined. Animals were first assayed for behavior and then mounted and imaged to quantify the number of RIM neurons. We excluded animals with 3 RIM-like cells and analyzed data for animals with either 2 RIMs (complete rescue of the RIM lineage cell death) or 4 RIM-like cells (no rescue of RIM lineage cell death) (Figure 3C). Based on rates of inheritance of the array in both RIM sisters, we estimated that in the 2-RIM animals, the other undead neuron types that express the *inx-19* promoter would retain the rescuing array at the following frequencies: ASI, 0.26; ADL, 0.22; ASK, 0.18; ADA, 0.26; PHB, 0.26; PVQ, 0.30; RIC, 0.51 (Methods). Of these, only ASI is implicated in reversals^21^. Thus, 2-RIM animals have the same number of RIM neurons as the wild type, but otherwise have a nervous system that is largely similar to that of *ced-3* mutants, with minor stochastic differences mostly outside the circuit for reversals.

We found that mosaic animals with two RIM neurons had significantly more and shorter reversals compared to *ced-3* animals and compared to 4-RIM-like mosaics (2-RIM = 4.1 reversals/min; *ced-3* = 3.1 reversals/min; p = 0.0172; 4-RIM = 2.9 reversals/min; p = 0.004) (Figure 3D). Reversals were also shorter in duration for *ced-3* mutant worms with only two RIM neurons (Figure 3E). The major difference between these groups is the number of undead RIM-like neurons: all three groups have large numbers of other undead cells. Together, these data indicate that the number of RIM-like neurons is a primary determinant of reversal frequency and duration. We note that animals with two RIMs were not identical to the wild type: although wild-type and 2-RIM animals had the same number of RIMs, the 2-RIM animals exhibited a modest decrease in reversals (wild type = 5.4 reversals/min; *ced-3* animals with 2 RIMs= 4.1 reversals/min; p=0.001) (Figure 3D and 3E). These data suggest that besides the undead RIM-like neurons, other undead neurons also contribute to the behavioral difference observed for *ced-3* animals. Overall, we conclude that undead RIM sisters are true neurons that exhibit coordinated activity, functionally integrate into the reversal circuit, and alter a behavior key to *C. elegans* navigation.

### Lack of programmed cell death causes additional specific behavioral alterations

To further investigate the functional consequences of undead neurons, we performed a series of assessments of PCD mutants in different behavioral states. We first performed a closed-loop high-throughput behavior assay in the absence of food to assess various aspects of locomotion while in the ‘local search’ phase of foraging^21,37–39^. Consistent with our manual analysis, we found that PCD mutants initiate fewer reversals (Figure 4A) and have longer reversals (Figure 4B). We tested two alleles each of *ced-3* and *ced-4* and found that all four mutants differed from wild type, with the alleles *ced-3(n3692)* and *ced-4(n1162)* causing slightly stronger phenotypes than *ced-3(n717)* and *ced-4(n3141)*. We also observed that all four mutants spent an amount of time in a paused state similar to that of the wild type (Figure S3A) but had altered turning behavior (Figure S3B). As omega turns coupled to reversals are tuned by the RIM circuit^40^, we further evaluated this behavior. The frequency of reversals coupled to omega turns (reversal omegas) and pure reversals were both decreased in PCD mutant worms (Figure S3C and S3D). Interestingly, there was a greater reduction in reversal omegas by *ced-4* mutants, with the fraction of reversals being coupled to omega turns significantly lower in *ced-4* mutants compared to *ced-3* mutants (Figure 4C, S3C). This observation suggests differing effects of *ced-3* and *ced-4* mutations on the deaths of cells that influence turning behavior. Also, PCD mutants have decreased forward velocity compared to wild-type animals, while reverse velocity is unchanged (Figure 4D, S3E). This observation indicates that other aspects of locomotion in addition to reversal behavior is altered in PCD mutants.

**Figure 4.**
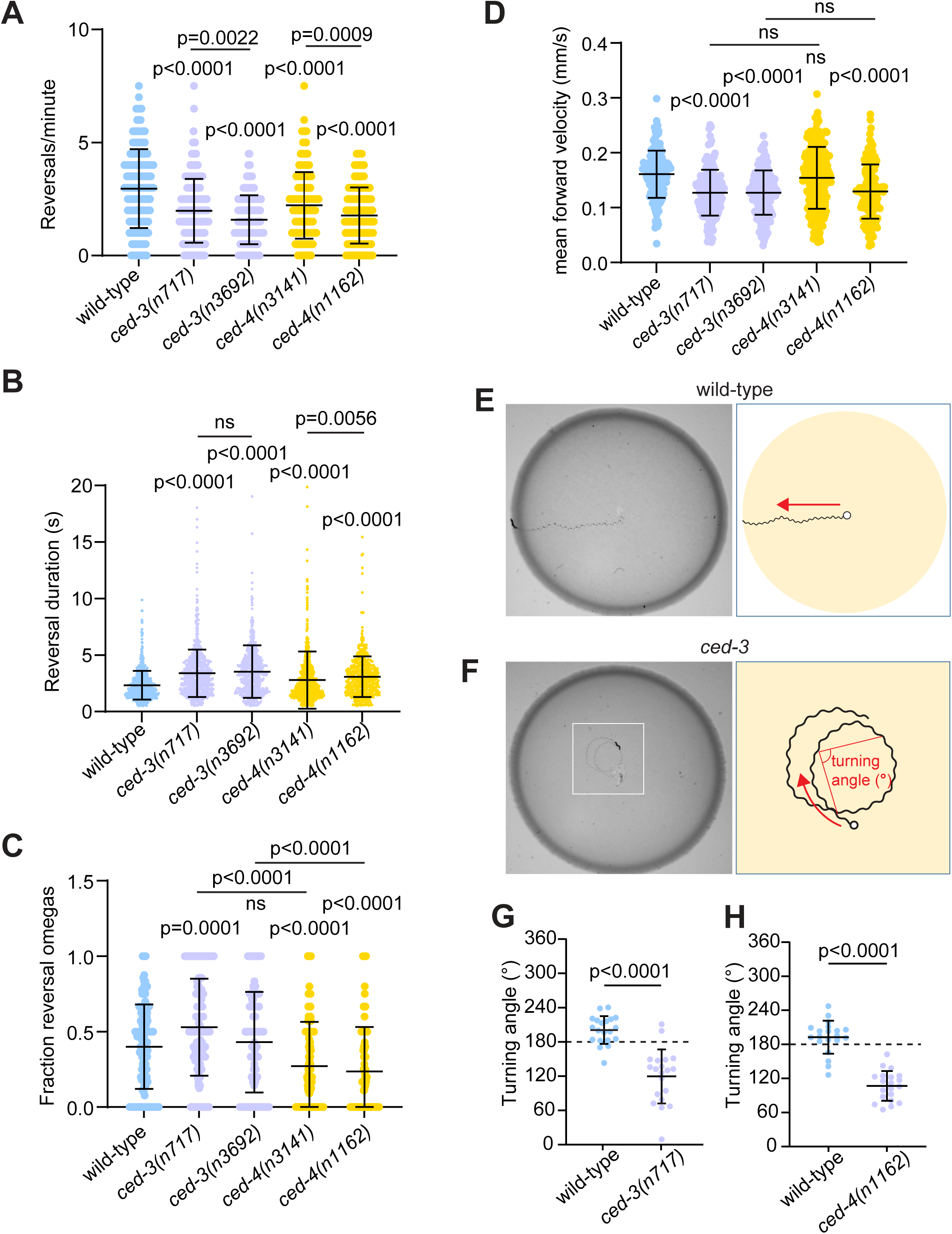
Analysis of cell-death mutant behavior. (**A-D**) Quantification of behavior in a high-throughput analysis off-food. Data are mean ± S.D. with individual data points shown. P-values directly above data represent comparison with wild-type (wild-type, n=158; *ced-3(n717),* n=194, *ced-3(n3692)*, n=187, *ced-4(n3141*), n=249, *ced-4(n1162),* n=163). (A) Cell-death mutants reverse less frequently in a high-throughput analysis of behavior off food. (B) Reversal duration is increased in cell death mutants. (C) The fraction of reversals coupled to omega turns (reversal omegas) is reduced in *ced-4*, but not *ced-3* mutants. (D) Forward velocity is reduced in cell-death mutants, except *ced-4(n3141)*. (**E-F**) Worm tracks on bacterial lawn. A wild-type worm moves straight (E), while a *ced-3(n717)* knockout worm tends to turn (F). (**G-H**) Quantification of the turning angle between control and *ced-3(n717)* (G) and between control and *ced-4(n1162)* (H). Worm strains used in panel H contain *nIs177[Pceh-28::4xNLS::GFP].* Data are mean ± S.D with n=20. Statistical significance was determined by Welch’s t-test.

We further examined forward locomotory movement of *ced-3* mutant animals in the presence of food. We observed that *ced-3* animals tended to turn toward their dorsal side while moving forward on a bacterial lawn, whereas wild-type animals showed no dorsal or ventral bias in their movement (Figure 4E and 4F). To quantify this difference, we measured the dorsal-side turning angles of worm tracks on a bacterial lawn (Figure 4F). Turning angle was quantified as the angle between two successive movement steps (five sinusoidal waves), such that the angle would be 180 degrees for animals moving in a straight line, < 180 degrees for animals turning dorsally, and >180 degrees for animals turning ventrally (Methods). We found that the wild-type turning angle averaged close to 180 degrees, displaying no strong dorsal or ventral bias. By contrast, *ced-3(n717)* mutants had significantly smaller turning angles than wild-type animals, indicating a bias toward dorsal turning (Figure 4G). Another mutant defective in PCD — *ced-4(n1162)* — similarly showed smaller dorsal turning angles than the wild type (Figure 4H), indicating that this behavioral abnormality is caused by a defect in PCD.

To more broadly explore the landscape of behavioral abnormalities of *ced-3* animals, we performed automated high-resolution analysis of freely-behaving *ced-3* animals in the presence of food. This unbiased analysis revealed additional significant differences. The rates of pharyngeal pumping and of defecation were significantly different between *ced-3* animals and the wild type, although the magnitude of these differences was small (Figure S3F and S3G).

The rates of egg laying were similar on average, but *ced-3* animals exhibited significantly more variability than the wild type (Figure S3H). By contrast, forward and reverse velocities were not significantly different (Figure S3I and S3J). Overall, our analyses support the idea that undead neurons modulate specific behaviors in specific ways, rather than causing global effects on all aspects of nervous system function.

## Discussion

We have demonstrated that PCD plays a critical role in controlling the circuits and computations of the *C. elegans* nervous system. We showed that specific undead cells, the sisters of the RIM neurons, can become functional neurons that are wired into a stereotyped circuit and influence behavior. These data suggest that changes in PCD on evolutionary timescales could support the tuning of nervous system composition to achieve desired computational outputs.

The two RIM neurons are elements of a circuit in the worm brain that computes the critical decision choice between two opposite modes of worm behavior: forward and backward movement. Each RIM neuron has a sister that is normally removed by PCD. In the absence of PCD, these RIM-proximate undead cells become RIM-like neurons and are integrated into the RIM circuit based on four criteria: (1) they expressed a RIM fate marker; (2) they made synapses with RIM’s normal presynaptic partners, the AIB neurons; (3) they displayed calcium activity that is time-locked with reversals and indistinguishable from the activity of the RIMs themselves; and (4) they modified reversal behavior.

Our data indicate that the activity of the undead RIM-like neurons resulted in fewer transitions from forward to backward movement and an increased duration of backward movement. The RIM interneurons modulate the transitions between forward and reverse locomotion, incorporating factors involving both internal state and external stimuli^21,37,38,40–43^. RIM drives different outcomes in locomotion behavior depending on its activity and its mode of transmission^21,22,26,40–42^. When depolarized, RIM stabilizes reversals through chemical synapses that lead to longer reversals, by inhibiting the transition to forward locomotion promoted by AVB neurons^44^ and possibly also through depolarization of the backward command AVA neurons^20,40,45^. RIM neurons can also suppress reversal initiation through electrical synapses from RIM to AVA when RIM is hyperpolarized, increasing forward run duration^40^. The behavioral alterations driven by the additional RIM-like neurons is likely a result of additional chemical synapses further stabilizing reversals and extending their length, ultimately resulting in fewer reversal events. As we see an increase in reversal duration, which is not altered by knockdown of *unc-9* gap junctions^40^, electrical synapses are unlikely to be the major mediator of behavioral alterations driven by additional RIM-like neurons. We found that turning behavior varied between *ced-3* and *ced-4* mutants, with reversal omegas less frequently executed by *ced-4* mutants. This difference might be driven by differences in the cell deaths that are blocked in these mutants or by a difference in non-apoptotic functions of *ced-3* and *ced-4*. *ced-4* regulates cell size^46^ and *ced-3* regulates developmental gene expression^47^, either or both of which could alter behavior through neuronal development. Interestingly, our data indicate that the removal of the undead RIM-like neurons did not completely revert reversal behavior to normal patterns, suggesting that other undead neurons can also affect this computation.

Failure of PCD dramatically alters the neuronal composition of *C. elegans*. The cell lineage contains 94 cell deaths that have neurons as their closest relatives. In the cases we examined, neuronal-proximate undead cells usually became neuron-like, expressing terminal differentiation markers and adopting morphologies similar to those of their lineage-proximate neuronal relatives. In some cases undead cells can adopt the neuronal fate of a sublineage homolog rather than of their closest relative, as has been observed for the P cell lineages that give rise to the VC motor neurons and, in PCD mutants, their undead VC homologs^14,48^. We additionally found that the undead sisters of the RIM neurons became functional neurons that can be wired into a stereotyped circuit and alter intrinsic behavior. These results raise the possibility that all 94 neuronal-proximate undead cells can become functional elements of the nematode nervous system, representing an increase in total neurons of 31% and possibly causing massive alterations in the normally stereotyped neuronal wiring and function. Although we describe some specific novel behavioral alterations of *ced-3* mutants, it seems likely that other aspects of behavior are perturbed in these mutants. It is possible that in some circuits a homeostasis mechanism compensates for the presence of extra neurons to maintain normal behavior.

A key function of nervous systems is to generate highly specific intrinsic behaviors. These intrinsic behaviors can be tuned over evolutionary timescales by alterations to the genome, with selection for changes that confer advantage. Because undead cells can differentiate into neurons, it has been proposed that cell death can alter neuronal circuits^6,18,19^. Our data indicate that cell death in neuronal cell lineages provides a reservoir for latent circuits. Either physiological or evolutionary alterations in cell death might access these latent circuits and drive novel behaviors. Furthermore, because in many cases the fates of undead cells are variable, so might be the resulting behavioral changes—conceptually similar to other phenotype variations arising from biological noise^49–52^, but with a mechanism that also involves altered information processing by neural circuits. Over time, by altering the genetic control of PCD, evolution could sculpt behaviors for maximum fitness in changing environments. Comparative analysis of the nematodes *C. elegans* and *P*. *pacificus* has shown that variations in cell-death patterning alter nervous system composition, which alongside evolutionary changes to neurite morphology and synaptic connections can shape nervous system function^53^. The human brain also computes intrinsic behaviors, and cell death is a prominent feature of mammalian brain development^54^.

Our results suggest that abnormalities in PCD in humans might lead to altered intrinsic behaviors, possibly manifesting as neuropsychiatric disease or abnormalities in motor control.

## Acknowledgments

We thank members of the Hammarlund laboratory for comments and suggestions. We thank the Caenorhabditis Genetic Center (funded by National Institutes of Health (NIH) Office of Research Infrastructure Programs P40 OD010440) for *C. elegans* strains. M.H. acknowledges the Whitman and Fellows program at MBL for providing funding and space for discussions valuable to this work.

## Author contributions

Conceptualization: ADK, MH, DL, HRH Methodology: ADK, MMT, SK, DL, SB, TS

Investigation: ADK, MMT, SK, DL, SB, TS, NK, CEE, NVP

Visualization: ADK, MMT, DL, SB

Funding acquisition: DCR, SF, HRH, AML, VV, MH Supervision: DCR, SF, HRH, AML, VV, MH Writing – original draft: ADK, MH

Writing – review & editing: ADK, MMT, SK, DL, SB, TS, NK, CEE, NVP, DCR, SF, HRH, AML, VV, MH

## Data and materials availability

All data are available in the manuscript or the supplementary materials.

## Methods

### EXPERIMENTAL MODEL AND STUDY PARTICIPANT DETAILS

#### C. elegans strains

All materials generated for this work are available from the authors. *C. elegans* strains (genotypes in table S2) were maintained at 20°C on NGM plates seeded with OP50 *E. coli* according to standard methods. The wild-type reference strain was N2 Bristol. Some strains were provided by the *Caenorhabditis* Genetics Center (CGC).

Molecular biology and transgenic lines

Plasmids were assembled using Gateway recombination (Invitrogen) or Gibson Assembly (New England Biolabs). For a list of plasmids and primers used see Table S3. Detailed cloning information will be provided upon request. Transgenic worms were created by microinjection of plasmids as previously described^55^ with co-injection markers *odr-1p::RFP* or *unc-122p::RFP.* Integrations were performed using UV-TMP as previously described^56^.

### METHOD DETAILS

#### Labeling and fluorescence microscopy of undead cells, neuronal fate markers

Undead cells were labeled by GFP-H2B fusion protein driven by 2788 bp of the *egl-1* promoter, 5820 bp 3’ UTR and includes the *egl-1* intron. For imaging, animals were immobilized with 25 mM levamisole (Sigma) and mounted on a 3% agarose pad on a microscope slide. Images were acquired as 0.2 µm Z-stacks using a Leica DMi8 microscope with a Hamamatsu Flash Orca 4.0 camera and VT-iSIM system (BioVision) and Metamorph software (Molecular Devices) with 40X HC Plan Apo NA 1.3 oil, 63X HC Plan Apo NA 1.4 oil, and 100X HC Plan Apo NA 1.47 oil objectives (Leica). Image analysis was performed using FIJI^57^. For PVD dendrites, the total number of quaternary dendrites were quantified 200 µm anterior of the most posterior PVD cell body.

#### RIM-specific expression

The genomic sequence between the *gcy-13* and F23H12.7 genes has previously been shown to drive strong and specific expression in RIM^23^. To drive expression in RIM neurons, we amplified a 2 kb segment between *gcy-13* and F23H12.7 (see Table S4 for primer sequences). We validated this driver by comparing its expression to previously-reported RIM-specific GFP expression^23^. We refer to plasmids containing this sequence as ‘pRIM’.

#### Labeling, microscopy and scoring of AIB-RIM synapses in wild-type and *ced-3(n717)* worms

AIB presynapses were labeled by GFP-tagged RAB-3 driven by the *inx-1* promoter, RIM postsynapses were labeled with tagRFP-tagged GLR-1 driven by the pRIM promoter, and RIM was labeled with BFP driven by the *cex-1* promoter. Animals were immobilized with 25 mM levamisole and mounted on a 3% agarose pad on a slide. Images were acquired as 0.2 µm Z-stacks using an iSIM microscope (BioVision) with a 100x oil objective and Metamorph software (Molecular Devices). We created a maximum intensity Z-projection of the upper 50% of the Z-stack to visualize only one side of the worm. The distal segment of the neurites was then straightened computationally. Z-projected images of the AIB::RAB-3::GFP channel were scored for patterning of normal synapses. Images for which there was increased synapse distribution and/or synaptic varicosities were scored as abnormal.

#### Calcium-imaging microscopy hardware

We used a modular confocal imaging system with a 60x, 1.3-NA silicone immersion objective (Olympus). To ensure slight movements with no displacement outside of imaging field, worms were semi-immobilized by placing them on a thin 1 x 1-cm 7% agar pad covered with a 22 x 22-mm #1.5 coverslip. 2-3 μL M9 buffer were added between the agar pad and the coverslip to increase surface tension, which further improved the immobilization. To obtain dual-color images, fluorescent proteins (mCherry and GCaMP6s) in RIM neurons were simultaneously excited by 561 nm and 488 nm laser lines (Andor ILE) to uniformly illuminate a 200 x 200-μm field of view. The emission light was projected onto the sensors of two sCMOS cameras (q) using a spinning disk confocal setup (Andor Dragonfly 200 Series), a 565 nm long-pass dichroic mirror and mCherry/GFP band-pass filters. To perform volumetric imaging, we mounted the objective on a nano positioning piezo (Pizeoconcept FOCHS.100). The full 4.2 megapixel camera sensor was used with 4 x 4 binning applied, resulting in images of 512 x 512 pixels with a 0.4 μm pixel size.

During acquisition, a DAQ device (Measurement Computing USB-1208HS-4A) was programmed to provide the primary clock to synchronize exposure times of cameras and the motion of the piezo. Camera exposure reset and the piezo moved to a new Z value once every two clock cycles. The piezo visited 25 different Z values spaced at 1.2 μm, resulting in volumes consisting of 25 stacks taken at 4 Hz. Data acquisition and image analysis were managed by custom software written in Python, which was run in a Windows 10 environment.

#### Extracting calcium traces

To extract signals, first we annotated all RIM neurons in each frame throughout the recordings. Next, pixel values in a 11 x 11 x 5-pixel region around each annotated neuron were sorted, and the mean of the first 40 pixel values was computed. Signal in each channel was modeled by 3 multiplying terms plus a time-independent noise term: The first term corresponded to the real fluorescent signal, the second term was a decaying function of time used to model the photobleaching effect, and the third term corresponded to artifacts shared between the red channel and the green channel. The fluorescent protein responsible for the signal in the red channel (mCherry) was calcium-insensitive, which meant the first term in our model was a constant; thus, a double exponential function was fitted to estimate the product of the first and the second terms, and the 0.8 quantile of all pixels was used to estimate the time-independent noise. Next the third term was computed and was used to remove artifacts from the green channel. In the same way, noise was subtracted from the green channel. To correct for the photobleaching effect in the green channel, only timepoints that were not within an event were used to fit a double exponential function, and the magnitude was proportionally adjusted to normalize traces after applying the correction.

#### Averaging reversals

For each RIM neuron, 6-second time windows of GCaMP6s signals around the reversal initiations were cropped, and the average value from t=-3 s to t=0 s was used as the baseline to calculate the signal change ratio. The change ratio was averaged across all reversal initiations and was plotted as a black line. The grey area corresponds to the standard error of the mean.

#### Correlation coefficient

For each animal, pairwise Pearson Correlation Coefficients between left-right neurons and wild-type-undead neurons were calculated and averaged between all animals. p-values were calculated and revealed no significant differences between calculated average values, which indicate strong wild-type-undead correlation.

#### Spaghetti plot

We used the time points at which reversal motions were initiated, and cropped a 10 s time window from 3 s before up to 7 s after these time points from each time series. In these cropped time series, we took the average of the first 3 s, and used it as a baseline (F0) for normalization.

#### Reversal Assay

The reversal assay was performed as previously described^58^. Young adults were transferred to unseeded NGM plates for 1 min before being recorded for 2 min. Any change from forward to backward movement was counted as a reversal during the 2-min interval. Reversal length was annotated using BORIS (Behavioral Observation Research Interactive Software)^59^.

#### Mosaic Analysis of Behavior

For RIM-specific CED-3A rescue in *ced-3(n717)* mutants, worms carrying the *pinx-19::CED-3A* array, indicated by the *punc-122::RFP* co-injection marker (expressed in coelomocytes), were selected and assayed for reversals, then mounted and imaged to determine the number of RIM and RIM-like neurons according to the previously described microscopic techniques.

In these experiments, the number of RIMs was experimentally determined. However, the *pinx-19::CED-3A* transgene is expressed not only in RIM and RIM-like neurons but also in other cell types^35,36^. If the array segregates into another cell type in which the *inx-19* promoter is active and that also has an undead relative, the array will also restore cell death there. To estimate this background, we calculated the stability of the array.

Out of 181 animals assayed (all of which expressed the coelomocyte marker, so all of which carried the array in at least some cells), 39 had two RIM neurons. This result indicates that the *pinx-19::CED-3A* array segregated through all cell divisions into the RIM sisters, where it rescued the *ced-3* mutation, thus allowing these cells to die. The only cell the RIM neurons and the coelomocytes have in common is the single-cell embryonic P0 cell. To segregate into both RIM sisters, the array must segregate through two common cell divisions (into ABp) and then through 8 cell divisions into the left RIM sister and 8 cell divisions into the right RIM sister. Thus, the array must segregate successfully through 18 cell divisions to yield 2 RIMs. Since the frequency of 2-RIM animals was 39/181 = 0.21547, the average probability of successful array transmission at each of these 18 mitotic cell divisions was the 18^th^ root of 0.21547, or 0.9182.

Using this probability, we can estimate the fraction of animals with two RIMs that also retain the array in other cells where it might function: that is, other cells in which the *inx-19* promoter is active and that also have an undead relative. There are 8 neuron types that fit this description; all are pairs of bilaterally-symmetric cells. Because we know that the array is present in the entire RIM lineage in these animals, we can calculate the fraction of animals that retain the array in these other lineages based on the average probability of successful array transmission at each mitotic division, assuming similar mosaicism rates across the lineage. This fraction will depend on the number of divisions each cell has after it diverges from the RIM lineage. We calculate the following fractions: ASI, 0.26; ADL, 0.22; ASK, 0.18; ADA, 0.26; PHB, 0.26; PVQ, 0.30; RIC, 0.51. Similarly, based on the array’s being present in at least the P0 cell, we can calculate the fraction of animals that retain the array in these other lineages in 4-RIM animals, which have lost the array somewhere in both the RIM left and right lineages: ASI, 0.22; ADL, 0.19; ASK, 0.15; ADA, 0.22; PHB, 0.22; PVQ, 0.25; RIC, 0.43.

#### Behavioral Data Collection and Analysis: Off Food

The wild-type, *ced-3(n717)*, *ced-3(n3692), ced-4(n3141),* and *ced-4(n1162)*, strains used in this study were grown at 20℃ on a standard nematode growth media (NGM) plate with OP50 (*E. coli*) as a food source. On the day of experiment, 6-7, day 1 young adult worms were transferred to an unseeded NGM plate (100 mm x 15 mm, Lab Express, Cat. # 5001-100). The animals were allowed to acclimatize to the new plate for 1 minute before collecting behavior measurements for 2 minutes using the previously described closed-loop high throughput behavior assay setup^60,61^. Briefly, the system captured images at 30 frames per second and a behavior classification algorithm was used to track and determine various behavior parameters for each of the worm as described in detail in a previous work^60^. Custom MATLAB scripts were written to quantify various behavior parameters as previously described^62^. A lot of times, worms went out of the field of view, or collided with another worm, or entered calibration zone^60^. Those worms were not included in the analysis. Only those worms which were continuously tracked for the whole duration of the assay (2 minutes) were included in the analysis.

#### Detection of reversal events

A reversal is defined as an instance when a worm’s velocity dips below -0.02 mm/s. The total number of reversals over the period of 2 minutes of the behavior assay was determined to calculate reversals/minute. The inbuilt MATLAB function ’findpeaks’ was used to detect reversals. The ’minpeakdistance’ parameter for this function was set to be 1.5 seconds to prevent over counting reversals which are part of the same reversal bout.

#### Determination of reversal duration

Reversal duration is the amount of time for which the worm’s velocity was below the threshold of -0.02 mm/s in each reversal bout.

#### Detection of turning events

Turn is defined as an instance when worm’s ellipse ratio dips below 3.1^61^. These are omega turns. The total number of turns over the period of 2 min behavior assay was determined to calculate turns/minute.

#### Determination of fraction of time spent in pause state

Pause state is defined as the instances when the worm’s velocity is between -0.02 mm/s and +0.02 mm/s. The total number of frames in pause state is determined for each worm to estimate fraction of time paused

#### Turning Angle

Five sinusoidal waves were defined as one movement step. The angle between two successive movement steps was defined as the turning angle. Single animals were transferred onto an OP50-seeded Petri plate and removed after1 – 2 minutes. Track images were captured and turning angles were measured using ImageJ.

#### Behavioral Data Collection and Analysis: On Food

L4 animals were picked approximately 24 hr before the start of the assay. All assay plates were seeded the night before with 200 µL of *E. coli* OP50 and allowed to dry overnight at room temperature. The assay plates were standard 10 cm diameter petri dishes with low-peptone (0.4 g/L) NGM. All behavioral recordings were conducted using a custom-built closed-loop tracking system at 20 Hz frequency and at 1.4 um/pixel resolution, as previously described^63^. On the day of the assay, adult animals were singly picked onto the assay plates, and plates were taped onto the microscope stage face down without a lid and on top of spacers to prevent condensation. After a 15 min period of acclimatization, assay animals were tracked for 3-6 hr, and behavioral parameters (locomotion, egg-laying, feeding and defecation) were extracted using a custom-built R software suit as previously described^63^.

### QUANTIFICATION AND STATISTICAL ANALYSIS

For Figure 3A, 3B, 3D, 3E, 4-D, 4G, 4H, S3A-S3J Unpaired two-tailed Welch’s t tests were used. For Figure S1J, unpaired two-tailed Student’s t tests were used. For categorical data, two-sided Fisher’s exact tests were used. Analysis was performed using Prism 9. Data is represented as mean and standard deviations (SD). Asterisks representing p values are as follows: *p<.05, **p<.01, ***p<.001, ****p<.0001.

---

**Figure S1.**
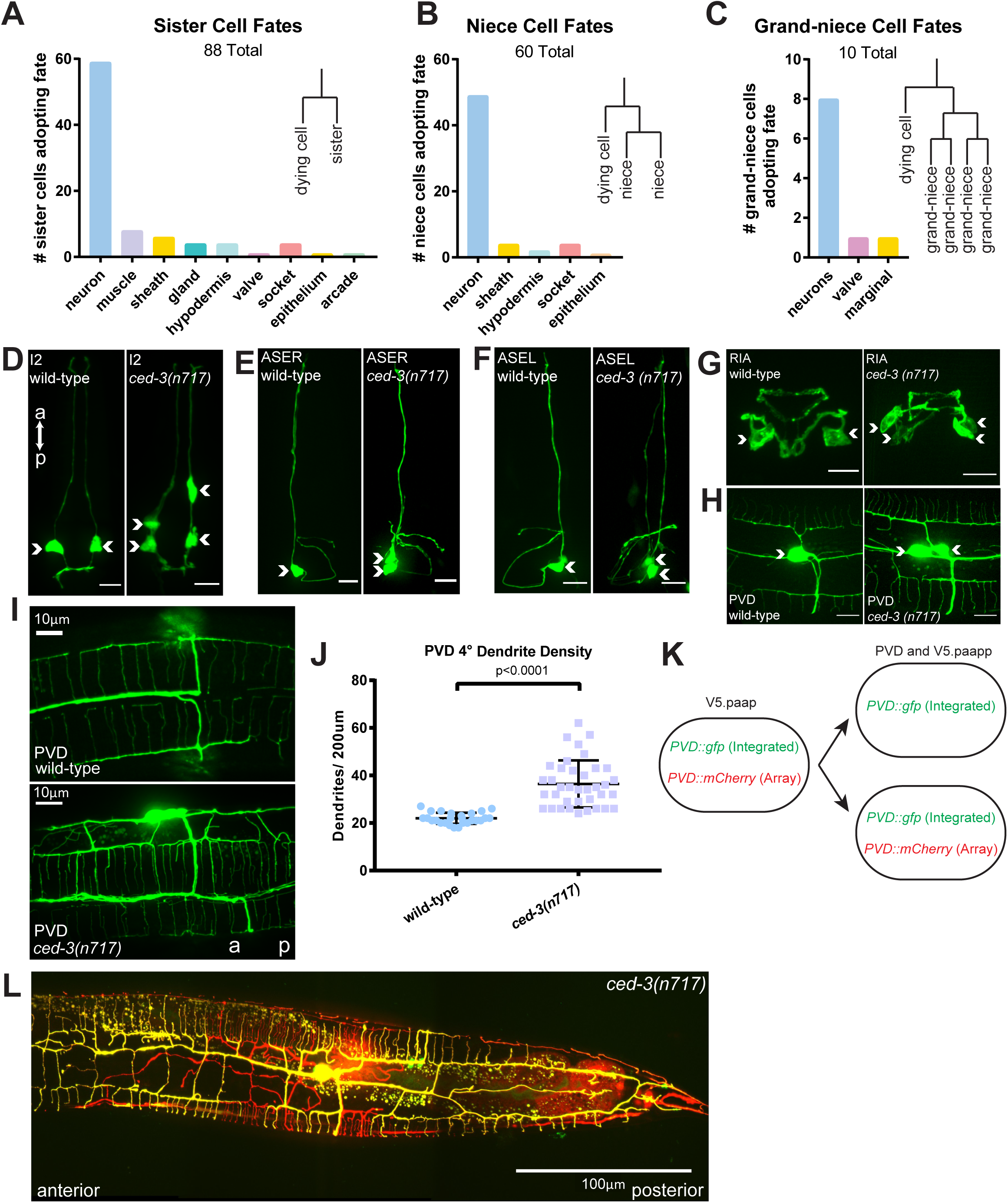
In *ced-3* mutants, undead cells adopt fates similar to those of their sister cells. (**A-C**) Fates and archetypical lineages of terminally differentiated relatives for all 131 cells removed by PCD in *C. elegans* hermaphrodites. (**D-H**) I2, ASER, ASEL, RIA, and PVD neurons each have sister cells that die in wild-type worms. In *ced-3(n717)* worms, additional cells express GFP markers for each of these neuronal fates (white arrows). 10 μm scale bars. (**I**) PVD quaternary dendrites viewed on the ventral side of the worm. (**J**) Quantification of quaternary dendrite density in wild-type and *ced-3(n717)* mutants. Data are mean ± S.D. with individual data points shown (wild type, n=26; *ced-3(n717),* n=37). (**K**) Schematic of mosaic PVD labeling. A GFP label is integrated into the genome and inherited by both wild-type and undead PVD cells. An mCherry PVD label is expressed as an extrachromosomal array, which in some animals is inherited by only one of the two PVDs on one side (either the wild-type or the undead cell). This allows visualization of the dendrites from each cell. (**L**) Mosaic PVD labeling demonstrates that both wild-type and undead PVDs produce dendrites. Statistical significance was determined by Student’s t-test.

**Figure S2.**
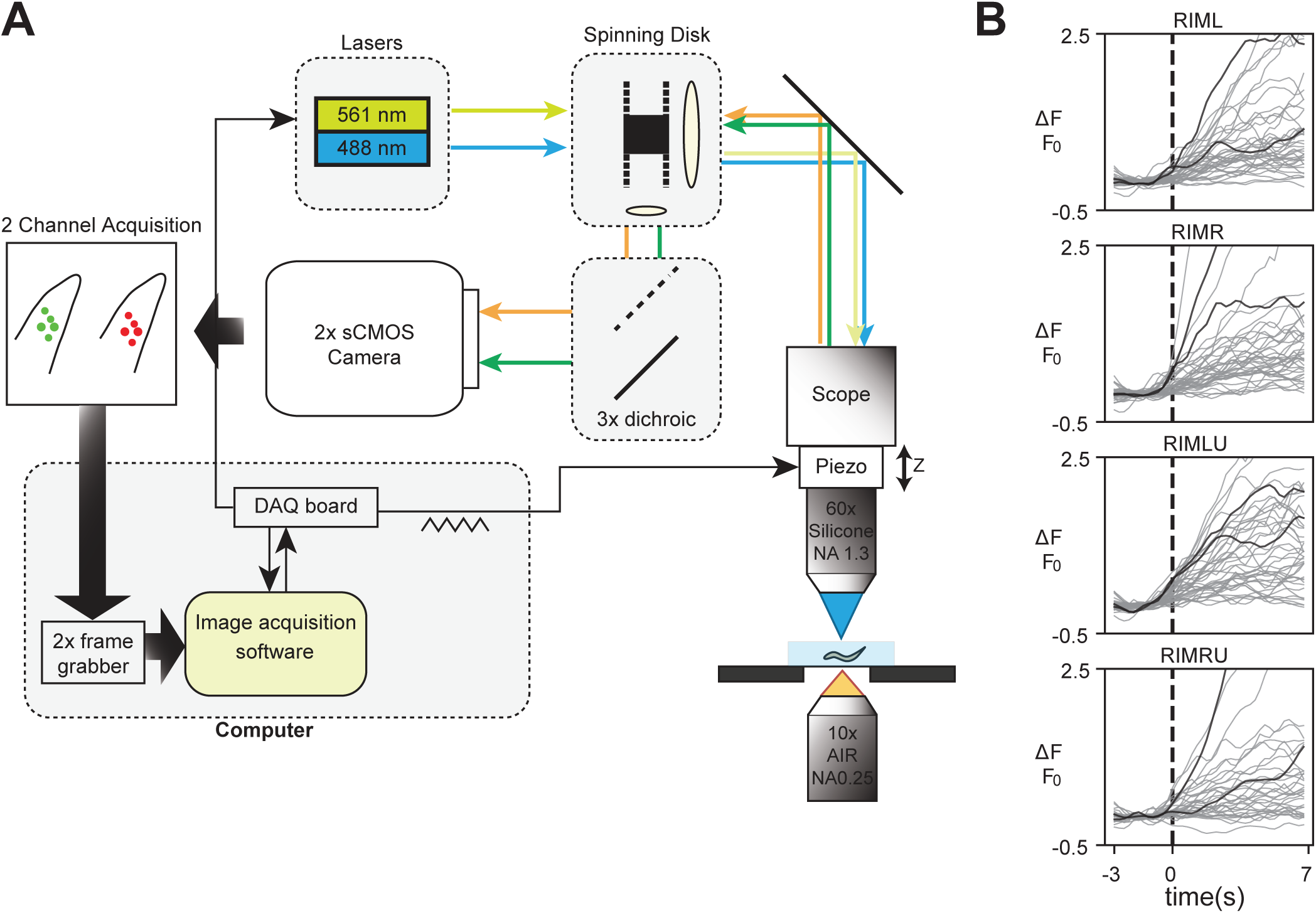
Function of undead RIM neurons. (**A**) Schematic of calcium imaging. (**B**) Spaghetti plot of all individual traces for average data shown in Figure 2E. Traces in black are two separate reversal events shown in Figure 2C.

**Figure S3.**
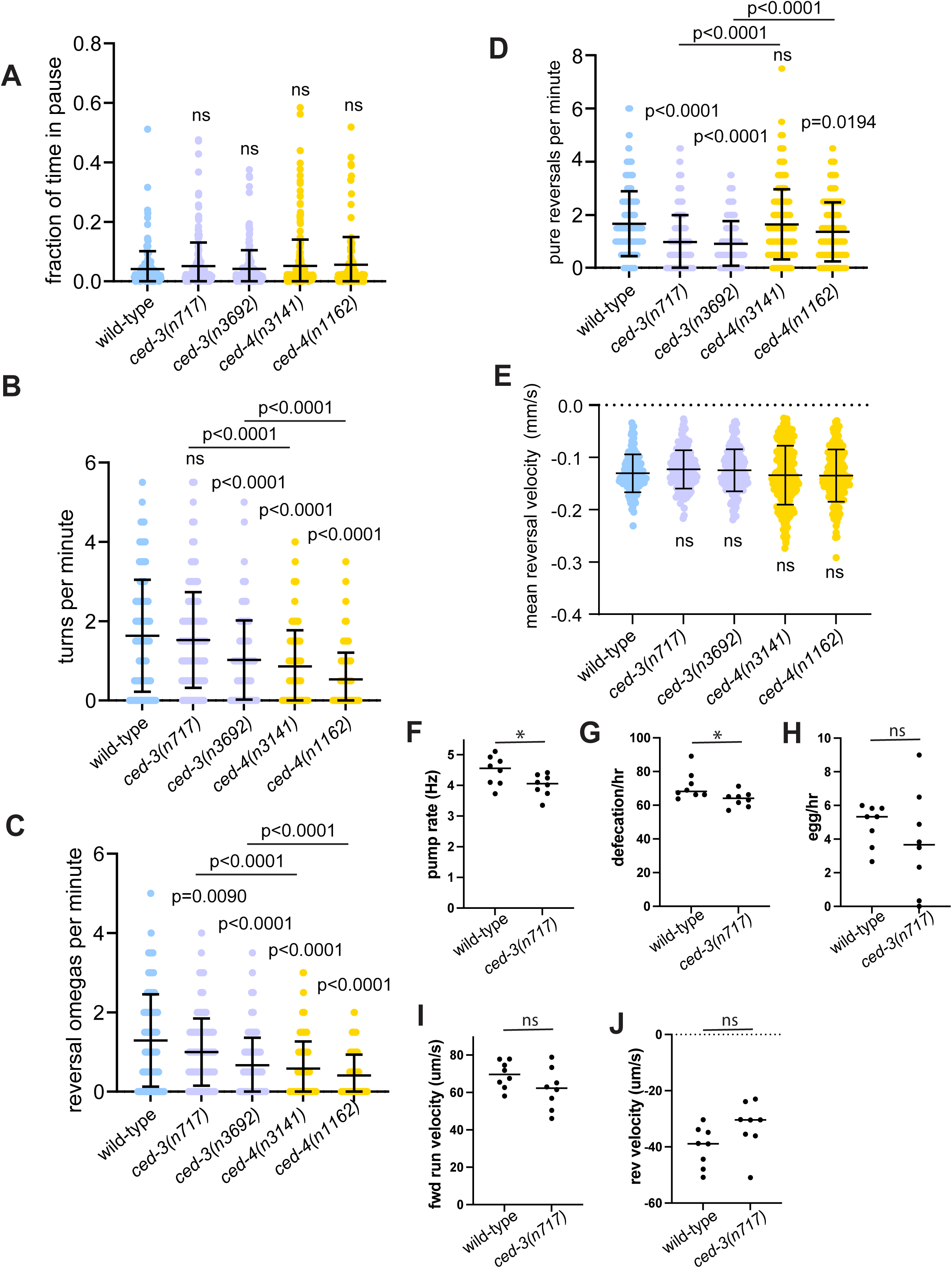
Additional analysis of *ced-3* behavior. (**A**) Cell-death mutants spend a similar fraction of time in a paused state compared to wild-type. (**B**) Cell-death mutants, except for *ced-3(n717)* initiate fewer turns than wild-type, with *ced-4* animals initiating less than *ced-3.* (**C**) Cell-death mutants initiate fewer reversals coupled to omega turns. (**D**) *ced-3* mutants initiate fewer pure reversals than wild-type and *ced-4* mutants. (**E**) Mean reversal velocity is unchanged in cell-death mutants. (**F**) *ced-3(n717)* worms have lower pharyngeal pumping rates than wild-type worms. (**G**) Defecation rates are lowered in *ced-3(n717)* animals. (**H**) Egg-laying rate is more variable among individuals for *ced-3* mutant animals. (**I,J**) Forward and reverse run velocity is not significantly different between wild-type and *ced-3(n717)* animals. Statistical significance was determined by Welch’s t-test.

