## Supplemental Tables for "Programmed Cell Death Modifies Neural Circuits and Tunes Intrinsic Behavior"

| **Table S1. Fates of terminally differentiated relatives of undead cells** | | | | | | | | |
| --- | --- | --- | --- | --- | --- | --- | --- | --- |
| **Undead Cell** | **Differentiated Cell Relation** | **Sister Cell** | **Niece 1** | **Niece 2** | **Grand-niece 1** | **Grand-niece 2** | **Grand-niece 3** | **Grand-niece 4** |
| **ABalaaaalar** | sister | AINL |  |  |  |  |  |  |
| **ABalaaaalpa** | sister | ILshL |  |  |  |  |  |  |
| **ABalaaaarla** | sister | RMEL |  |  |  |  |  |  |
| **ABalaaaarra** | sister | RMER |  |  |  |  |  |  |
| **ABalaaaparl** | sister | ILshDL |  |  |  |  |  |  |
| **ABalaaapplr** | sister | ILshDR |  |  |  |  |  |  |
| **ABalaapaaal** | sister | AINR |  |  |  |  |  |  |
| **ABalaapaapa** | sister | ILshR |  |  |  |  |  |  |
| **ABalaapapa** | nieces |  | CEPsoVR | dead |  |  |  |  |
| **ABalaapappa** | sister | CEPsoVR |  |  |  |  |  |  |
| **ABalaappaa** | nieces |  | AVAR | OLQsoVR |  |  |  |  |
| **ABalaapppap** | sister | RIAR |  |  |  |  |  |  |
| **ABalaappppap** | sister | IL1R |  |  |  |  |  |  |
| **ABalapaaaap** | sister | AVHL |  |  |  |  |  |  |
| **ABalapaapap** | sister | RIAL |  |  |  |  |  |  |
| **ABalapaappap** | sister | IL1L |  |  |  |  |  |  |
| **ABalapapaa** | niece + grandnieces |  | URYDL |  | OLQDL | dead |  |  |
| **ABalapapapap** | sister | OLQDL |  |  |  |  |  |  |
| **ABalapappaap** | sister | IL1DL |  |  |  |  |  |  |
| **ABalappaaa** | nieces |  | RID | dead |  |  |  |  |
| **ABalappaapp** | sister | RID |  |  |  |  |  |  |
| **ABalappapap** | sister | AVHR |  |  |  |  |  |  |
| **ABalapppapap** | sister | OLQDR |  |  |  |  |  |  |
| **ABalappppaap** | sister | IL1DR |  |  |  |  |  |  |
| **ABalpaaaapp** | sister | m1VL |  |  |  |  |  |  |
| **ABalpaaapap** | sister | m2L |  |  |  |  |  |  |
| **ABalpaaappa** | sister | MCL |  |  |  |  |  |  |
| **ABalpapaapp** | sister | arc ant V |  |  |  |  |  |  |
| **ABalpapapapa** | sister | SMBDL |  |  |  |  |  |  |
| **ABalpapappa** | sister | SMBVL |  |  |  |  |  |  |
| **ABalpappaaa** | nieces |  | I2L | dead |  |  |  |  |
| **ABalpappaapp** | sister | I2L |  |  |  |  |  |  |
| **ABalpapppap** | sister | I1L |  |  |  |  |  |  |
| **ABalppaaaa** | nieces |  | AVAL | OLQsoVL |  |  |  |  |
| **ABalppaapa** | nieces |  | CEPsoVL | dead |  |  |  |  |
| **ABalppaappa** | sister | CEPsoVL |  |  |  |  |  |  |
| **ABalppapppap** | sister | IL1VL |  |  |  |  |  |  |
| **ABalpppaapv** | sister | OLLshL |  |  |  |  |  |  |
| **ABalpppappa** | nieces |  | ASKL | dead |  |  |  |  |
| **ABalpppapppp** | sister | ASKL |  |  |  |  |  |  |
| **ABalppppaav** | sister | ADLL |  |  |  |  |  |  |
| **ABalppppapp** | nieces |  | OLLL | dead |  |  |  |  |
| **ABalppppapap** | sister | OLLL |  |  |  |  |  |  |
| **ABalppppppap** | sister | ASEL |  |  |  |  |  |  |
| **ABaraaaapp** | nieces |  | e1D | dead |  |  |  |  |
| **ABaraaaapaa** | sister | e1D |  |  |  |  |  |  |
| **ABaraapapaad** | sister | NSML |  |  |  |  |  |  |
| **ABaraapppaad** | sister | NSMR |  |  |  |  |  |  |
| **ABarapaaapp** | sister | m1VR |  |  |  |  |  |  |
| **ABarapaapap** | sister | m2R |  |  |  |  |  |  |
| **ABarapapaaa** | nieces |  | I2R | dead |  |  |  |  |
| **ABarapapaapp** | sister | I2R |  |  |  |  |  |  |
| **ABarapappap** | sister | I1R |  |  |  |  |  |  |
| **ABarappaaap** | sister | RIPR |  |  |  |  |  |  |
| **ABarappaapp** | sister | hyp1V |  |  |  |  |  |  |
| **ABarappapapa** | sister | SMBDR |  |  |  |  |  |  |
| **ABarapppppap** | sister | IL1VR |  |  |  |  |  |  |
| **ABarpaaapp** | nieces |  | OLQshDR | CEPshDR |  |  |  |  |
| **CEMDR** | sister | URADR |  |  |  |  |  |  |
| **ABarpapaapa** | nieces |  | CEPDR | URXR |  |  |  |  |
| **CEMDL** | sister | URADL |  |  |  |  |  |  |
| **ABplaaaaapa** | nieces |  | CEPDL | URXL |  |  |  |  |
| **ABplaapapppp** | sister | ASIL |  |  |  |  |  |  |
| **ABplapaaaaa** | nieces |  | ADEL | ADAL |  |  |  |  |
| **P1aap** | nieces |  | AVFL | VB1 |  |  |  |  |
| **QLaa** | sister | PQR |  |  |  |  |  |  |
| **Table S1 continued** | | | | | | | | |
| **Undead Cell** | **Differentiated Cell Relation** | **Sister Cell** | **Niece 1** | **Niece 2** | **Grand-niece 1** | **Grand-niece 2** | **Grand-niece 3** | **Grand-niece 4** |
| **QLpp** | nieces |  | PVM | SDQL |  |  |  |  |
| **V5Lpaapp** | sister | PVDL |  |  |  |  |  |  |
| **P9aap** | nieces |  | VA9 | VB10 |  |  |  |  |
| **P11aap** | nieces |  | VA11 | dead |  |  |  |  |
| **P11aaap** | sister | VA11 |  |  |  |  |  |  |
| **ABplapapppa** | niece + grandnieces |  | dead |  | PLML | ALNL |  |  |
| **ABplapappppp** | nieces |  | PLML | ALNL |  |  |  |  |
| **ABplapppaap** | sister | PVQL |  |  |  |  |  |  |
| **ABplapppapa** | nieces |  | HSNL | PHBL |  |  |  |  |
| **TLpppp** | nieces |  | PHCL | PLNL |  |  |  |  |
| **ABplpaaappap** | sister | OLQVL |  |  |  |  |  |  |
| **CEMVL** | sister | AMsoL |  |  |  |  |  |  |
| **ABplpaappppp** | sister | CEPVL |  |  |  |  |  |  |
| **ABplpappap** | nieces |  | RMEV | exc cell |  |  |  |  |
| **ABplppaaap** | niece + grandnieces |  | SIBDL |  | RICL | dead |  |  |
| **ABplppaaaapa** | sister | RICL |  |  |  |  |  |  |
| **ABplppaapaa** | sister | RIML |  |  |  |  |  |  |
| **ABplpppapp** | nieces |  | PHshL | hyp8 |  |  |  |  |
| tail spike | sister | hyp10 |  |  |  |  |  |  |
| **ABpraaaaapv** | sister | OLLshR |  |  |  |  |  |  |
| **ABpraaaappa** | nieces |  | ASKR | dead |  |  |  |  |
| **ABpraaaapppp** | sister | ASKR |  |  |  |  |  |  |
| **ABpraaapaav** | sister | ADLR |  |  |  |  |  |  |
| **ABpraaapapp** | nieces |  | OLLR | dead |  |  |  |  |
| **ABpraaapapap** | sister | OLLR |  |  |  |  |  |  |
| **ABpraaapppap** | sister | ASER |  |  |  |  |  |  |
| **ABpraapapppp** | sister | ASIR |  |  |  |  |  |  |
| **ABprapaaaaa** | nieces |  | ADER | ADAR |  |  |  |  |
| **Wap** | nieces |  | AVFR | VB2 |  |  |  |  |
| **P2aap** | nieces |  | VA2 | VB3 |  |  |  |  |
| **QRaa** | sister | AQR |  |  |  |  |  |  |
| **QRpp** | nieces |  | AVM | SDQR |  |  |  |  |
| **V5Rpaapp** | sister | PVDR |  |  |  |  |  |  |
| P10aap | nieces |  | VA10 | VB11 |  |  |  |  |
| **P12aap** | nieces |  | VA12 | dead |  |  |  |  |
| **P12aaap** | sister | VA12 |  |  |  |  |  |  |
| **P12pp** | sister | hyp12 |  |  |  |  |  |  |
| **ABprapapppa** | niece + grandnieces |  | dead |  | PLMR | ALNR |  |  |
| **ABprapappppp** | nieces |  | PLMR | ALNR |  |  |  |  |
| **ABprapppaap** | sister | PVQR |  |  |  |  |  |  |
| **ABprapppapa** | nieces |  | HSNR | PHBR |  |  |  |  |
| **TRpppp** | nieces |  | PHCR | PLNR |  |  |  |  |
| **ABprpaaappap** | sister | OLQVR |  |  |  |  |  |  |
| **CEMVR** | sister | AMsoR |  |  |  |  |  |  |
| **ABprpaappppp** | sister | CEPVR |  |  |  |  |  |  |
| **ABprppaaap** | niece + grandnieces |  | SIBDR |  | RICR | dead |  |  |
| **ABprppaaaapa** | sister | RICR |  |  |  |  |  |  |
| **ABprppaapaa** | sister | RIMR |  |  |  |  |  |  |
| **ABprpppapp** | nieces |  | PHshR | hyp9 |  |  |  |  |
| tail spike | sister | hyp10 |  |  |  |  |  |  |
| **MSaaaappa** | sister | m7D |  |  |  |  |  |  |
| **MSaaappa** | sister | vpi3D |  |  |  |  |  |  |
| **MSaapaapap** | sister | g1AL |  |  |  |  |  |  |
| **MSaapapap** | sister | g2L |  |  |  |  |  |  |
| **MSapaapa** | M lineage (muscle) |  |  |  |  |  |  |  |
| **MSappapa** | sister | mu bod |  |  |  |  |  |  |
| **MSpaaaaap** | sister | M4 |  |  |  |  |  |  |
| **MSpaapp** | grandnieces |  |  |  | M1 | dead | mc3DR | vpi1 |
| **MSpaapaap** | sister | M1 |  |  |  |  |  |  |
| **MSpapaapap** | sister | g1AR |  |  |  |  |  |  |
| **MSpapapap** | sister | g2R |  |  |  |  |  |  |
| **MSppaapa** | sister | mu int R |  |  |  |  |  |  |
| **MSpppaaa** | Z1 lineage |  |  |  |  |  |  |  |
| **MSpppapa** | sister | mu bod |  |  |  |  |  |  |
| **Caapap** | sister | DVC |  |  |  |  |  |  |

| **Table S2: *C. elegans* strains used in this study.** | |  | | |
| --- | --- | --- | --- | --- |
| **Strain Name** | **Strain Details** | | **Source** | **Details** |
| N2 | *C. elegans* wild isolate | | Caenorhabditis Genetics Center | WT |
| MT1522 | ced-3(n717) IV | | Caenorhabditis Genetics Center | cell death mutant |
| MT12054 | ced-3(n3692)IV | | H. Robert Horvitz Lab | cell death mutant |
| MT2547 | ced-4(n1162)III | | Caenorhabditis Genetics Center | cell death mutant |
| MT10034 | ced-4(n3141)III | | H. Robert Horvitz Lab | cell death mutant |
| IK718 | njIs12[glr-3p::glr-1::GFP+glr-1p::RFP+ges-1p::RFP] | | Caenorhabditis Genetics Center | RIA label |
| XE2016 | njIs12[glr-3p::glr-1::GFP + glr-1p::RFP + ges-1p::RFP]; ced-3(n717) IV | | This paper | cell death mutant, RIA label |
| NY2045 | ynIs45[flp-15p::GFP] I; him-5(e1490) V | | Caenorhabditis Genetics Center | I2 label |
| XE2019 | ynIs45[flp-15p::GFP]; ced-3(n717)IV | | This paper | cell death mutant, I2 label |
| NC1687 | wdIs52[F49H12.4p::GFP + unc-119 (+)] II | | Caenorhabditis Genetics Center | PVD label |
| XE1894 | wdIs52[F49H12.4p::GFP + unc-119 (+)] II; ced-3(n717) IV | | This paper | cell death mutant, PVD label |
| XE2212 | wdIs52[PVD GFP (F49H12.4p::GFP), unc-119(+)] II wpIs119[F49H12.4p::NLS_mCherry::unc-54UTR] | | This paper | cytoplasmic GFP and nuclear mCherry in PVDs |
| XE2213 | wdIs52[PVD GFP (F49H12.4p::GFP), unc-119(+)] II ced-3(n717) IV wpIs119[F49H12.4p::NLS_mCherry::unc-54UTR] | | This paper | cell death mutant, cytoplasmic GFP and nuclear mCherry in PVDs |
| OH3191 | otIs3[gcy-7p::GFP + lin-15(+)] V | | Caenorhabditis Genetics Center | ASEL label |
| XE1801 | ced-3(n717) IV; otIs3[gcy-7p::GFP + lin-15(+)] V | | This paper | cell death mutant, ASEL label |
| OH3192 | ntIs1[gcy-5p::GFP + lin-15(+)]V | | Caenorhabditis Genetics Center | ASER label |
| XE1802 | ced-3(n717) IV; ntIs1[gcy-5p::GFP + lin-15(+)]V | | This paper | cell death mutant, ASER label |
| LX836 | lin-15(n765ts )X; vsIs44 [tph-1p::GFP] V | | Michael R. Koelle Lab | NSM label |
| XE2001 | ced-3(n717) IV; vsIs44 [tph-1p::GFP] V | | This paper | cell death mutant, NSM label |
| PHX2910 | sbIs2910[gcy13p::GCaMP6s + gcy-13p::tagRFP] | | This paper | RIM label |
| XE2718 | ced-3(n717) IV sbIs2910[gcy13p::GCaMP6s + gcy-13p::tagRFP] | | This paper | cell death mutant, RIM label |
| XE2738 | ced-3(n717) IV sbIs2910[gcy13p::GCaMP6s + gcy-13p::tagRFP] olaEx5053[inx-19p::ced-3a + cex-1p::bfp + unc-122p::rfp] | | This paper | cell death mutant, RIM-specific cell death rescue, RIM label |
| XE2739 | wpEx474[inx-1p::rab-3::gfp+ cex-1p::bfp + gcy-13p::glr-1::tagRFP] | | This paper | AIB-RIM synaptic label |
| XE2851 | ced-3(n717); wpIs132[egl-1p::gfp::h2b::egl-1UTR +Podr-1::rfp] X; wpEx491[gcy13p::tagRFP] | | This paper | cell death mutant, undead cell label with RIM tagRFP label |
| XE2740 | ced-3(n717) IV wpEx474[inx-1p::rab-3::gfp+ cex-1p::bfp + gcy-13p::glr-1::tagRFP] | | This paper | cell death mutant, AIB-RIM synaptic label |
| XE2844 | ynIs45[flp-15p::GFP] I; ced-3(n717); wpEx489[tph-1p::RFP + unc-122p::RFP] | | This paper | cell death mutant, I2 label, NSM label |
| XE2845 | ces-2(n732ts); vsIs44[tph-1p::GFP] V | | This paper | NSM lineage cell death mutant, NSM label |
| XE2197 | ced-3 (n717); wpIs132[egl-1p::gfp::h2b::egl-1UTR +odr-1p::rfp] X | | This paper | cell death mutant, undead cell label |
| XE2198 | wpIs132[egl-1p::gfp::h2b::egl-1UTR +odr-1p::rfp] X | | This paper | cell death WT, undead cell label |
| DCR5516 | olaIs67 [inx-1p(1 kb)::EGFP::Rab-3 + inx-1p(451bp)::mCherry + unc-122p::RFP];X | | *Sengupta T*. et al. 2021 | AIB synapse label |
| DCR5695 | ced-3(ola338); olaIs67 [inx-1p(1 kb)::EGFP::Rab-3 + inx-1p(451bp)::mCherry + unc-122p::RFP];X | | This paper | cell death mutant, AIB synapse label |
| DCR4894 | olaex2887 [inx-1p(1 kb)::EGFP::Rab-3 + inx-1p(1 kb)::mCherry::PHD + unc-122p::RFP] | | *Sengupta T*. et al. 2021 | AIB synapse label |

| **Table S3: DNA plasmids used in this study.** | | | |  |  |  |
| --- | --- | --- | --- | --- | --- | --- |
| **Plasmid name** | **Promoter** | **Size (bp)** | **Oligonucleotide 1 (5 → 3)** | **Oligonucleotide 2 (3 → 5)** | **Full Construct** | **Vector** |
| pAK025 | egl-1 | 2756 | gctgaaagagtgttgacagtaacc | gacatctccctactatctttcccctata | pegl-1::GFP:H2B::egl-1 3' UTR | pCFJ150 |
| pAK0031 | pF49H12.4 | 2025 | gtaagtgagggcaaagtgtgcatc | tcacaacgcacaggtttcac | pF49H12.4::NLS mCherry::unc-54 3' UTR | pDEST R4-R3 |
| pAK0024 | pF49H12.4 | 2025 | gtaagtgagggcaaagtgtgcatc | tcacaacgcacaggtttcac | pF49H12.4::mCherry::unc-54 3' UTR | pDEST R4-R3 |
| pAK0044 | gcy-13 | 2000 | aaaaattgctaaaagatttataaatcagcaagtggttt | ttgttatttgaaacttattacaaactttcaataatttttcaggac | pgcy-13::GCamP6s::unc-54 3' UTR | pCFJ150 |
| pAK0045 | gcy-13 | 2000 | aaaaattgctaaaagatttataaatcagcaagtggttt | ttgttatttgaaacttattacaaactttcaataatttttcaggac | pgcy-13::tagRFP::unc-54 3' UTR | pCFJ150 |
| pAK0047 | gcy-13 | 2000 | aaaaattgctaaaagatttataaatcagcaagtggttt | ttgttatttgaaacttattacaaactttcaataatttttcaggac | pgcy-13::GLR-1::tagRFP::unc-54 3' UTR | pCFJ150 |
| pDACR2245 | inx-1(1 kb) | 1000 | attattttctgtgcttttcacaaatacac | tccggcggacaagaac | inx-1p(1kb)::EGFP::Rab-3::unc-54 3' UTR | pSM |
| pDACR1412 | inx-1(451bp) | 451 | tatagttcttcatcttcttttttttaatatcctc | tccggcggacaagaac | inx-1p(451bp)::mCherry::unc-54 3' UTR | pDEST[4-3] |
| pDACR2404 | inx-1(1 kb) | 1000 | attattttctgtgcttttcacaaatacac | tccggcggacaagaac | inx-1p(1 kb)::mCherry::PHD::unc-54 3' UTR | pDEST[4-3] |
| pDACR3149 | cex-1 | 947 | ttggaactttaaacgggtttttaaatg | tctagaaatgaacattccatggg | pcex-1::mtagBFP1::unc-54 3' UTR | pDEST[4-3] |
| pDACR3315 | inx-19 | 5625 | acgtaccgaagagatgtg | CAGCAGTTTCCCTGAATTAAA | inx-19p::ced-3A::unc-54 3' UTR | pDEST[4-3] |
| pDACR489 | tph-1 | 3124 | ggtggtcttcccgcttgcaat | gtttttaggtagcattgctctcttcaatcat | tph-1p::tagRFP::unc-54 3' UTR | pDEST[4-3] |
|  |  |  |  |  | H2B: Histone H2B used for chromosomal targeting |  |
|  |  |  |  |  | NLS: Nuclear localization signal |  |
|  |  |  |  |  | PHD: Pleckstrin homology domain used for membrane targeting |  |

| **Table S4: Oligonucleotides used in this study.** | | |
| --- | --- | --- |
| **Oligonucleotide** | **Source** | **Identifier** |
| egl-1_5’_fwd: 5’- gctgaaagagtgttgacagtaacc -3’ | This study | egl-1_5’_fwd |
| egl-1_5’_rev: 5’- gacatctccctactatctttcccctata-3’ | This study | egl-1_5’_rev |
| egl-1_3’_1_for: 5’- ttttcgatctctccgtctccaactc-3’ | This study | egl-1_3’_1_for |
| egl-1_3’_1_rev: 5’-gtgtcaattagtattgcgcgag -3’ | This study | egl-1_3’_1_rev |
| egl-1_3’_2_for: 5’- ccggaaatccagtgtcaattag-3’ | This study | egl-1_3’_2_for |
| egl-1_3’_2_rev:5’- TTTGAGCTGGAACACTCGAGACGTACGG-3’ | This study | egl-1_3’_2_rev |
| F49H12.4_for: 5’- gtaagtgagggcaaagtgtgcatc-3’ | This study | F49H12.4_for |
| F49H12.4_rev: 5’- tcacaacgcacaggtttcac-3’ | This study | F49H12.4_rev |
| gcy-13_for: 5’- aaaaattgctaaaagatttataaatcagcaagtgg-3’ | This study | gcy-13_for |
| gcy-13_rev: 5’- atttgaaacttattacaaactttcaataatttttcaggac-3’ | This study | gcy-13_rev |
